# Molecular evolution across developmental time reveals rapid divergence in early embryogenesis

**DOI:** 10.1101/518621

**Authors:** Asher D. Cutter, Rose H. Garrett, Stephanie Mark, Wei Wang, Lei Sun

## Abstract

Ontogenetic development hinges on the changes in gene expression in time and space within an organism, suggesting that the demands of ontogenetic growth can impose or reveal predictable pattern in the molecular evolution of genes expressed dynamically across development. Here we characterize co-expression modules of the *C. elegans* transcriptome, using a time series of 30 points from early-embryo to adult. By capturing the functional form of expression profiles with quantitative metrics, we find fastest evolution in the distinctive set of genes with transcript abundance that declines through development from a peak in young embryos. These genes are highly enriched for oogenic function (maternal provisioning), are non-randomly distributed in the genome, and correspond to a life stage especially prone to inviability in inter-species hybrids. These observations conflict with the “early conservation model” for the evolution of development, though expression-weighted sequence divergence analysis provides some support for the “hourglass model.” Genes in co-expression modules that peak toward adulthood also evolve fast, being hyper-enriched for roles in spermatogenesis, implicating a history of sexual selection and relaxation of selection on sperm as key factors driving rapid change to ontogenetically distinguishable co-expression modules of genes. We propose that these predictable trends of molecular evolution for dynamically-expressed genes across ontogeny predispose particular life stages, early embryogenesis in particular, to hybrid dysfunction in the speciation process.

**Impact Summary:** The development of an organism from a single-celled embryo to a reproductive adult depends on dynamic gene expression over developmental time, with natural selection capable of shaping the molecular evolution of those differentially-expressed genes in distinct ways. We quantitatively analyzed the dynamic transcriptome profiles across 30 timepoints in development for the nematode *C. elegans.* In addition to rapid evolution of adult-expressed genes with functional roles in sperm, we uncovered the unexpected result that the distinctive set of genes that evolve fastest are those with peak expression in young embryos, conflicting with some models of the evolution of development. The rapid molecular evolution of genes in early embryogenesis contrasts with the exceptional conservation of embryonic cell lineages between species, and corresponds to a developmental period that is especially sensitive to inviability in inter-species hybrid embryos. We propose that these predictable trends of molecular evolution for dynamically-expressed genes across development predispose particular life stages, early embryogenesis in particular, to hybrid dysfunction in the speciation process.

## Introduction

Ontogenetic development hinges on the changes in gene expression in time and space within an organism. The dynamic molecular networks that specify cell proliferation and differentiation together produce morphogenesis, going from a single-celled zygote to a reproductively mature adult. Evolution favors maximal reproductive success to shape those gene expression dynamics and the functional properties of the proteins they encode, with the strength of selection pressures recorded in their sequences. Therefore, the demands of ontogenetic growth ought to impose or reveal predictable pattern in the molecular evolution of genes expressed dynamically across development (Raff 1996; Kalinka and Tomancak 2012). The rules, if any, that govern the molecular evolution of development must integrate adaptive evolution within the cellular constraints to forming a whole organism in embryogenesis and the life history constraints on a whole organism to reproduce successfully. We can address these issues from the perspective of genetic controls (e.g. *cis-* and trans-regulation) or from spatio-temporal dynamics in the formation of the structures of a complete organism.

A physical, spatial perspective motivates one means of molecular evolutionary predictability in development: tissue-or cell-specificity of gene expression will narrow the breadth of expression in space and consequently narrow the potentially negative pleiotropic effects of changes to gene expression or protein function (Stern 2000; Carroll 2005; Haygood et al. 2010; He et al. 2012). This logic about the impact of pleiotropy mirrors arguments for the disproportionate role of *cis-* regulatory changes in adaptive divergence, relative to trans-regulatory and coding changes (Wray 2007; Carroll 2008; Stern and Orgogozo 2008; Wittkopp and Kalay 2012). For example, mammalian genes with greater tissue-specificity of expression evolve faster in coding sequence but slower in terms of expression change (Liao and Zhang 2006b).

Temporal specificity of gene expression provides a parallel dimension to spatial specificity that can restrict or exacerbate the potential for pleiotropic effects of change to gene regulation or protein structure. Similar to the argument for spatial extent of gene activity, narrower duration of expression in ontogeny ought to narrow the potential for negative pleiotropic effects of changes to a given gene. A counter-argument, however, points out the unidirectional nature of time: changes to early points in development can cascade through ontogeny with disproportionate force (Poe and Wake 2004; Irie and Kuratani 2014; Arthur 2015). Because most new mutations with fitness effects are deleterious (Keightley and Lynch 2003), this “early conservation” or “generative entrenchment” view predicts slower evolution of genes expressed earlier in embryonic development, as has been reported for mouse and zebrafish (Roux and Robinson-Rechavi 2008; Irie and Kuratani 2014). By contrast, the most famous temporal paradigm derives from embryological observations of a ‘phylotypic stage’ with greatest phenotypic constraint relative to earlier and later timepoints in development, the ‘hourglass model’ (Raff 1996; Kalinka and Tomancak 2012). Applications of this idea to molecular data have renewed interest in it beyond morphology for diverse taxa, including *C. elegans* (Castillo-Davis and Hartl 2002; Cutter and Ward 2005; Levin et al. 2012; Zalts and Yanai 2017) and other invertebrates (Davis et al. 2005; Cruickshank and Wade 2008; Kalinka et al. 2010; Gerstein et al. 2014; Levin et al. 2016; Liu and Robinson-Rechavi 2018b), vertebrates (Hazkani-Covo et al. 2005; Domazet-Loso and Tautz 2010; Irie and Kuratani 2011; Piasecka et al. 2013; Liu and Robinson-Rechavi 2018a), and even plants (Quint et al. 2012; Drost et al. 2015). Different still, population genetics arguments about weaker purifying selection on genes expressed by just one sex, like maternal-effect gene products deposited in eggs, predict disproportionately rapid evolution of such maternally-deposited genes involved in early-embryogenesis of zygotes (Cruickshank and Wade 2008).

These ‘evo-devo’ ideas, however, largely focus on embryogenesis, and do not explicitly incorporate the entirety of ontogeny over an organism’s life cycle (Kalinka and Tomancak 2012). Ideas from the evolution of aging and senescence, by contrast, consider late life (Flatt and Schmidt 2009). In particular, the mutation-accumulation theory of aging predicts more rapid evolution of genes expressed following the onset of reproductive maturity than for those expressed earlier because diminishing reproductive value following maturity weakens the ability of selection to eliminate mutations (Medawar 1952; Charlesworth 1993; Promislow and Tatar 1998; Partridge 2001). Genes with expression in just one sex also ought to experience weaker purifying selection than other genes, leading to faster protein evolution, because mutations would be exposed to selection in just half of the population (Cruickshank and Wade 2008). Sexual contests and mate choice drive rapid divergence in morphological ornaments and their genetic underpinnings (Swanson and Vacquier 2002; Ellegren and Parsch 2007), so sexual selection and sexual conflict also predicts faster evolution of sex-biased genes and of genes expressed late in life, to the extent that their development gets specified toward adulthood. The coding sequences of adult-expressed genes do tend to evolve faster than embryonic genes in a number of taxa (Cutter and Ward 2005; Davis et al. 2005; Artieri et al. 2009; Liu and Robinson-Rechavi 2018a).

*C. elegans* and related nematodes are well-known for their similarity in form (Haag et al. 2007), despite the long times since species separated from one another (Cutter 2008). Indeed, the embryonic cell lineage of different *Caenorhabditis* species is outwardly preserved to an astonishing degree (Zhao et al. 2008; Memar et al. 2018), albeit with some key differences in timing of developmental milestones (Levin et al. 2012). Upon hatching at the end of embryogenesis, *C. elegans* individuals comprise 558 cells, then growing to become adult hermaphrodites with 959 somatic cells total (Sulston et al. 1983). The similarity of form across species, however, masks substantial evolution of genetic interactions as revealed by pronounced embryonic mortality in interspecies hybrids (Baird et al. 1992; Baird and Seibert 2013; Bundus et al. 2015). Developmental system drift is thought to underlie evolutionary change to spindle movement in the first cell division of embryos (Riche et al. 2013; Farhadifar et al. 2015; Valfort et al. 2018). Experiments also demonstrate that morphological stasis and even conserved expression patterns mask profound cis-regulatory divergence of conserved coding genes (Barriere et al. 2012; Barrière and Ruvinsky 2014; Verster et al. 2014; Barkoulas et al. 2016). Molecular evolution analysis of genes expressed differentially across post-embryonic development from microarray data reported faster evolution of coding sequences associated with the onset of reproductive maturity, but little directional effect of timing in embryogenesis (Cutter and Ward 2005). These collective observations motivate characterization of molecular evolution for gene expression dynamics across the entirety of ontogeny to explain the paradox of morphological conservation and hybrid dysfunction.

Here we test for evo-devo patterns of molecular evolution by characterizing co-expression modules of the *C. elegans* transcriptome over the full course of development, using functional principle components analysis (FPCA) on a time series of 30 points from early embryo to adults (Gerstein et al. 2010; Gerstein et al. 2014). By coarse graining the functional form of these ontogenetic trajectories of gene expression, we capture quantitative metrics that reveal how developmental dynamics relate to rates of molecular evolution. We find predictable trends of molecular evolution across ontogeny that are most conspicuous when analyzing ontogenetically co-expressed sets of genes, with implications for the genetics of post-zygotic reproductive isolation in the speciation process.

## Methods

### Expression data source and primary processing

We obtained RNAseq transcriptome sequences as sam-format files (mapped to *C. elegans* reference genome version WS248) from the public modENCODE data repository (http://data.modencode.org) for the *C. elegans* developmental time series for early embryos, each larval stage and young adult hermaphrodites (Supplementary Table S1) (Gerstein et al. 2010; Gerstein et al. 2014). We quantified expression for each gene using featureCounts (Liao et al. 2014), based on exon annotations of WS248 (transposable element and pseudogene annotations were excluded; exons corresponding to all alternative splice forms of a given gene contributed to expression quantification for that gene). We then normalized expression counts following the log-counts per million method of (Law et al. 2014). Embryonic transcriptomes included a single biological replicate per timepoint whereas larval and young adult transcriptomes included duplicates; given the high correlation between duplicates (r > 0.95), we used the average log-normalized expression for each larval and adult timepoint for subsequent analyses. We restricted our analyses to those 19,711 genes with an expression level ≥1 read count per million (cpm) in at least one timepoint (Robinson et al. 2010). We recalculated the log-cpm values for this set of 19,711 genes to account for the slight change in library sizes after the filtering step.

### Co-expression clustering and expression quantification of modules

To uncover and identify distinct sets of gene expression patterns over time across the 19,711 genes in the *C. elegans* transcriptome (co-expression “modules”), we performed a functional principal components analysis (FPCA). FPCA is appropriate for longitudinal datasets that may be sampled irregularly, with dense or sparse sampling, or with noisy values (Yao et al. 2005; Hall et al. 2006; Madrigal et al. 2018), as for this transcriptome time series with just a single replicate per timepoint. First, we applied FPCA to the log-normalized gene expression data, using the “FPCA” function in the R package fdapace, observing the first two components to cumulatively explain ~92% of the total variation. We then used each gene’s FPC scores of the first two components as input for the clustering algorithm, implemented through the “FClust” function in R that uses a Gaussian Mixture Model approach based on EMCluster (W.C. Chen and R. Maitra, 2015, http://cran.r-project.org/package=EMCluster). We determined the optimal number of co-expression clusters or modules in our analysis to be *k* = 14, based on minimizing the Bayesian information criterion (BIC) value. We varied *k* between 2 and 20 and observed minimum ΔBIC = 11.4 occurring between *k* = 12 and *k* = 14. Visual inspection of expression trends affirmed the biological relevance of choosing *k* = 14 co-expression modules to represent the variation in expression profiles in the *C. elegans* transcriptome time series. Based on the outputs of the clustering algorithm, we assigned each gene to the module for which the gene has the highest membership probability.

To summarize quantitatively the dominant trends in expression over time for each co-expression module, we fit orthogonal cubic polynomial functions with time to log-normalized expression values, rescaled using the “poly_rescale” function in the polypoly R package (T. Mahr, 2017, https://cran.r-project.org/package=polypoly). To relate the co-expression modules to each other, we then performed hierarchical clustering on the module-wise cubic polynomial regression coefficients. The goal with this functional analysis was not statistical testing of model complexity (e.g. linear vs. quadratic), but to use the parameter values of a flexible functional form as a quantitative metric of expression profile shape that can be compared across co-expression modules and across genes. The parameters extracted from the cubic fits summarize the overall expression level (α), increasing or decreasing trends in expression across development (β_1_), the degree of concave versus convex expression dynamics over ontogeny (β_2_), and how S-shaped are the expression dynamics (β_3_). In order to obtain a finer-grained view of the temporal trends, we also performed a gene-level analysis, in which we fit an orthogonal cubic polynomial to each individual gene expression profile and extracted the corresponding parameters for analysis.

Finally, we classified genes according to expression pattern in the simplest of ways, by grouping genes according to which timepoint they showed peak expression across the time series.

### Enrichment analysis

To investigate trends of genomic organization for each co-expression module, we used contingency tables and χ^2^-test statistics to test for non-random distributions of genes for each of the 14 modules across each of the 6 chromosomes in the genome. To achieve this, we arranged the data in 84 individual two-way contingency tables, so that we could obtain χ^2^-test statistics on 1 degree of freedom to test for an association within each module-chromosome combination. We further investigated trends of genomic organization by looking within chromosomes, at enrichment within the arm and center regions of each chromosome, with arm vs. centre domains defined by recombination rate breakpoint positions given by (Rockman and Kruglyak 2009). MtDNA genes were excluded for these analyses, and p-values were adjusted for multiple testing using the Holm–Bonferroni method.

We conducted gene ontology (GO) and phenotype enrichment analysis (PEA) tests using the list of genes in each co-expression module as input into the WormBase Enrichment Analysis Suite (Angeles-Albores et al. 2016; Angeles-Albores et al. 2018), obtaining Benjamini-Hochberg false discovery rate corrected p-values (q-values) for statistical significance. By also cross-referencing genes with the analysis of (Tu et al. 2015), we used their determination of operon identity and calculations of coding sequence divergence between orthologs of *C. elegans* and *C. briggsae* to quantify molecular evolution of protein sequence as *K*_A_, the rate of non-synonymous site substitution per non-synonymous site. Because of the saturated synonymous-site substitution rates (*K*_S_), we focus on *K*_A_ as a metric of protein molecular evolution rather than *K*_A_/*K*_S_ (Cutter and Ward 2005). Finally, we cross-referenced the genes in the transcriptome time series with those identified by (Ortiz et al. 2014) to have sex-neutral, oogenic or spermatogenic enrichment of expression in their analysis of *C. elegans* transcriptomes from dissected gonads.

## Results

### Ontogenetic expression dynamics define stereotypical transcriptomic patterns

We used functional principle components analysis (FPCA) to define 14 co-expression modules that describe clusters of the 19,711 genes that get expressed across 30 timepoints from early embryo through young adult stages of hermaphrodite *C. elegans* (Figure 1), based on ModENCODE transcriptome profiling data (Supplementary Table S1) (Gerstein et al. 2010; Gerstein et al. 2014). To obtain quantitative metrics describing the shape of each co-expression module, we then fit a cubic function to the gene expression profiles of each of the 14 developmental time series (Figure 1). The parameter values extracted from the cubic fits capture the overall expression level (α), increasing or decreasing trends in expression across development (β_1_), the degree of concave versus convex expression dynamics over ontogeny (β_2_), and how S-shaped are the expression dynamics (β_3_). When we then fit the cubic functional form to each gene individually (Supplementary Figure S1, Supplementary Figure S2), discriminant analysis demonstrated that values for these four parameters could correctly determine the co-expression module identity for 92.9% of genes, indicating that parameters from gene-wise cubic function fits capture well the key distinguishing features of ontogenetic expression dynamics.

**Figure 1.**
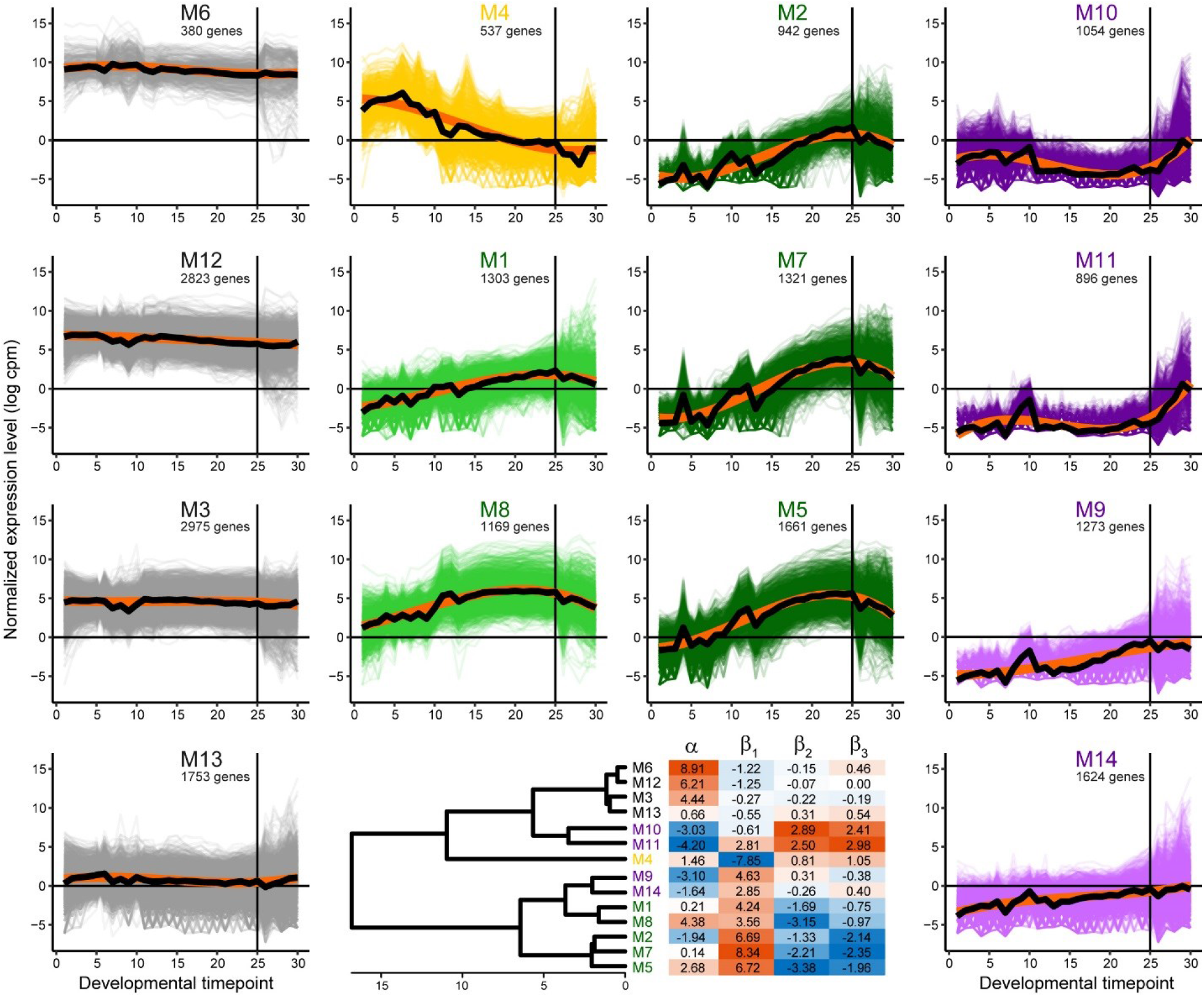
Ontogenetic time series of 19,711 *C. elegans* gene expression profiles clustered into 14 co-expression modules. Modules colored according to a trend of decreasing expression across development (yellow M4), peak expression in late embryogenesis (green M1, M8; M2, M7, M5), peak expression post-embryogenesis (purple M10, M11; M9, M14), or non-dynamic ‘constitutive’ expression across all 30 developmental timepoints (gray M6, M12, M3, M13). Thick black curves indicate expression trend across all genes in a module; thick orange curves indicate cubic polynomial fit to the expression trend. Similarity of module profiles indicated in dendrogram, with heatmap of parameter values from polynomial fit to each module expression trend (α = overall expression level, β_1_ = linear change over time, β_2_ = quadratic curvature, β_3_ = cubic S-shape to expression profile over development). Vertical line at developmental timepoint 25 indicates the end of embryonic development, followed by 5 post-embryonic timepoints; embryonic timepoints taken at 30 minute intervals, with 1 timepoint for each larval stage L1-L4 and young adult (Supplementary Figure S1) (Gerstein et al. 2010; Gerstein et al. 2014).

Four modules show consistent expression with little change across development (M3, M6, M12, M13). These ‘constitutive’ gene expression modules differ from one another primarily in the overall magnitude of expression (highest α=8.91 for M6, lowest α=0.66 for M13) and include the three largest modules by gene membership (M3, M12, M13) (Figure 1). By contrast, five modules exhibited hump-shaped expression dynamics with low expression in early embryos coupled to peak expression in late embryogenesis (β_1_>>0, β_2_<<0, β_3_<<0; M1, M2, M5, M7, M8). Module M4 was unique among all modules in showing peak expression in early embryogenesis, which then declined across developmental time (β_1_<<0). The four remaining modules displayed peak expression in post-embryonic stages (M9, M10, M11, M14), with especially strong up-regulation toward adulthood in M10 and M11 (Figure 1).

### Biased genomic architecture of ontogenetic gene expression modules

Upon defining these ontogenetically dynamic gene expression modules, we investigated their distinguishing features in terms of genomic organization, function and molecular evolution. Interestingly, genes from related expression profiles showed distinctive chromosome biases. Five modules were enriched on the X-chromosome, all of which corresponded to those with peak expression in late embryogenesis (M1, M2, M5, M7, M8; Figure 2). This genomic nonrandomness to expression co-variation in ontogeny suggests that chromatin regulation might influence the fitness effects of gene translocations in predictable ways. The early-embryogenesis module M4 showed the greatest chromosomal bias of any module, being >2-fold enriched on Chromosome II and tended to be underrepresented on all other chromosomes (Figure 2). Genes from those modules with peak post-embryonic expression, by contrast, showed enrichment on chromosomes IV and V (M9, M10, M11, M14), and highly-expressed ‘constitutive’ modules showed enrichment on chromosomes I and III (M3, M6, M12; Figure 2).

**Figure 2.**
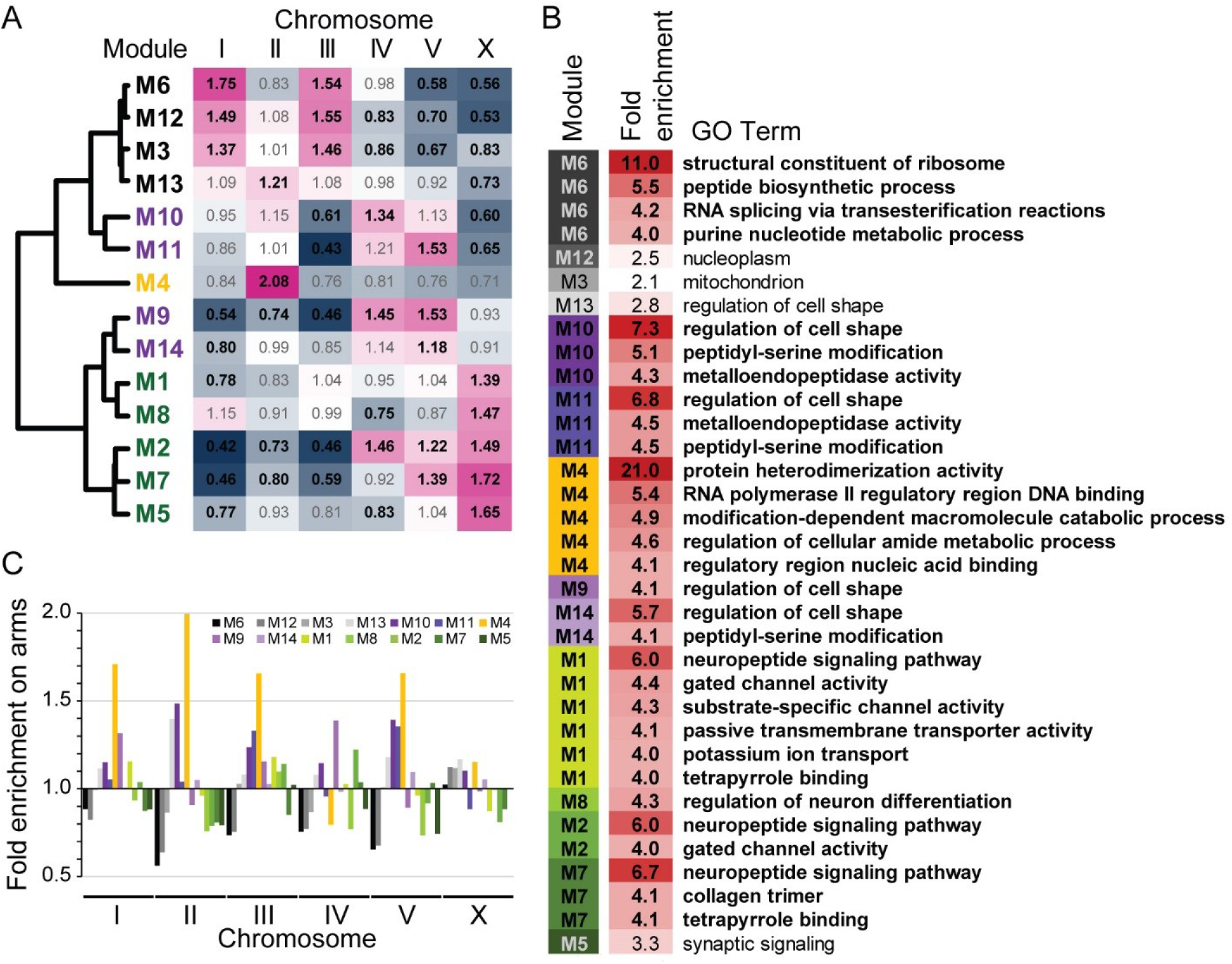
(A) Enrichment of gene membership among chromosomes for each co-expression module. Bold black text for observed/expected values in the heatmap indicates significant over-or under-enrichment (Holm-Bonferroni corrected p-values < 0.05,). (B) List of the 30 most enriched (>4-fold) gene ontology (GO) terms for each module, plus the single most enriched GO term observed for M3, M5, M12, M13 (all q-values < 0.005; 346 significantly enriched GO terms total across the 14 modules; Supplementary Table S1, Supplementary Table S2). (C) Enrichment of module gene membership on chromosome arms (values <1 imply enrichment in chromosome centers), where arm regions have higher recombination, higher density of repetitive elements, and lower gene density. Genome-wide significant enrichment on autosomal arms for M4, M10, and M13 and in centers for M5 and M12 (all Holm-Bonferroni corrected p-values < 0.003). Module identities colored and sorted by expression profile similarity as in Figure 1.

When we looked within chromosomes at their recombination domain structure of arms versus centers (Rockman and Kruglyak 2009), we found genes for most modules to be present in their expected proportions given chromosomal gene densities (Figure 2). However, genes in M4 were significantly enriched in arms on Chromosome II, the chromosome where M4 genes are exceptionally abundant, and also were elevated on arms relative to centers of other chromosomes (Figure 2). Post-embryonic modules M9 and M10, as well as the low-expression ‘constitutive’ module M13, also showed significant enrichment on arms of several chromosomes (Figure 2). By contrast, the highly-expressed ‘constitutive’ module M12 was under-enriched on the arms of Chromosomes II and V (Figure 2).

At a more local scale of genome organization, we found that three modules were hyper-enriched for membership in operons (Figure 3). Each of the highly-expressed ‘constitutive’ modules M3, M6 and M12 contain >40% of their genes in operons (Figure 3), compared to just 20.5% of coding genes overall occurring in operons. Of the remaining modules, only M13 (the fourth ‘constitutive’ module) and M8 had >10% operonic genes, and <4% of genes occurred in operons for all four modules with post-embryonic peak expression (M9, M10, M11, M14) (Figure 3).

**Figure 3.**
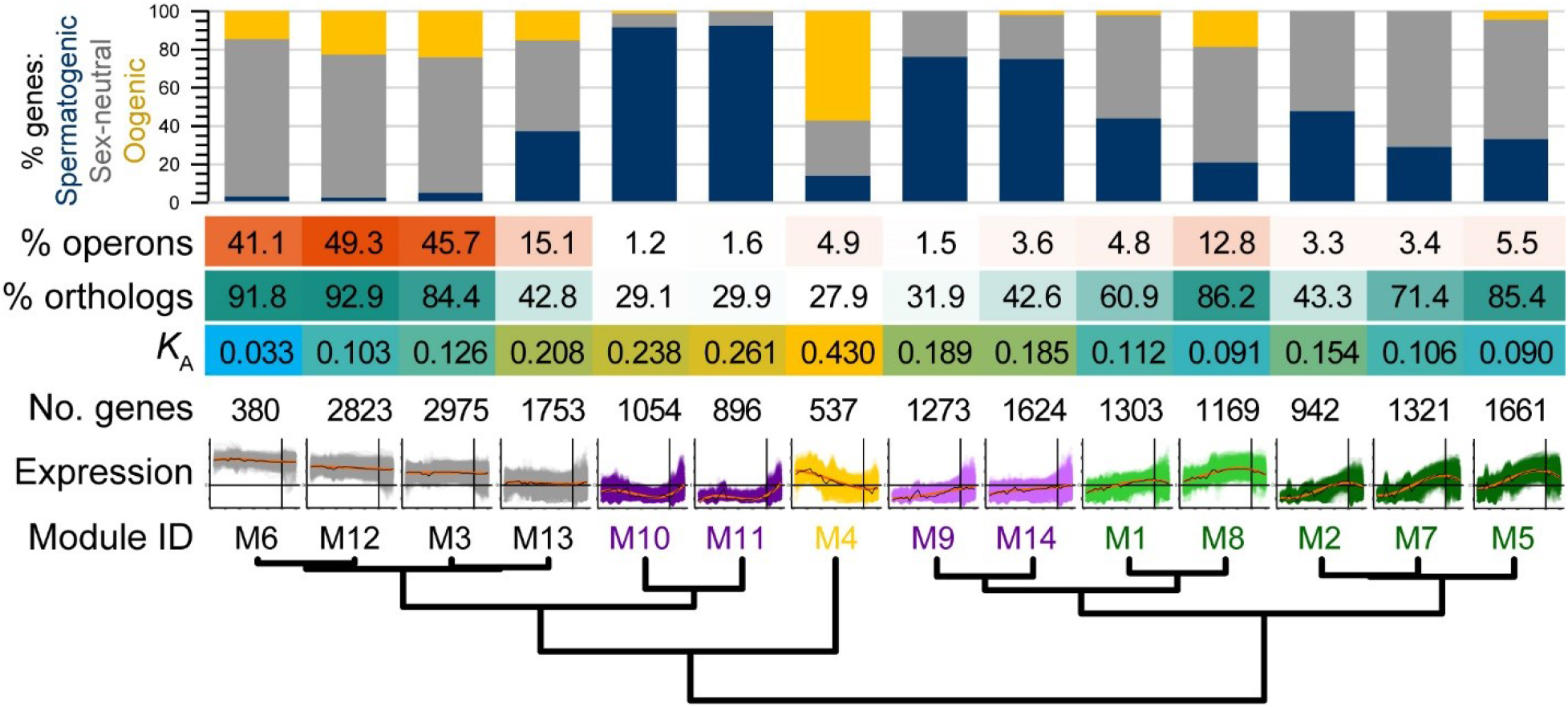
Functional and evolutionary properties of genes within each co-expression module. The proportion of genes with enrichment of spermatogenic, oogenic or sex-neutral expression categories defined by Ortiz et al. (2014), shown in the cumulative bar graph. Heat map shows the incidence of module genes in operons, the fraction of module members having orthologs in *C. briggsae,* and the median rate of non-synonymous site substitution (*K*_A_) as a measure of protein sequence divergence. Module order sorted by expression profile similarity as in Figure 1.

### Distinctive functional properties of ontogenetic gene expression modules

We cross-referenced the gene composition of co-expression modules with those gene sets identified by Ortiz et al. (Ortiz et al. 2014) to have sex-neutral, oogenic or spermatogenic enrichment of expression. These three expression categories had been inferred from differential expression of dissected gonads that had either active oocyte-only or sperm-only development (Ortiz et al. 2014). The early-embryogenesis module M4 showed extreme enrichment for oogenic genes (57%), with the next most enriched modules for oogenic genes being ‘constitutive’ modules M3 (24%) and M12 (23%) (Figure 3, yellow portion of bar plots). By contrast, the four modules with peak expression in post-embryonic stages contained almost no oogenic genes, instead being exceptionally enriched for spermatogenic genes (75% to 92%) (Figure 3; M9, M10, M11, M14). As expected of genes with sperm-related function (Reinke and Cutter 2009), operons were rarest in these modules (M9-M11, M14) (Figure 3). Eight of the 14 modules overall were comprised of >50% sex-neutral genes, including all five of those with peak expression late in embryogenesis, though three of the ‘constitutive’ modules contained the highest abundance of them (71% to 82%; M3, M6, M12) (Figure 3, gray portion of bar plots).

Gene ontology (GO) and phenotype enrichment analysis (PEA) further showed that the highly expressed ‘constitutive’ modules are enriched for basic cellular processes, like ribosomal and mitochondrial activity, embryonic defects, and chromosome segregation (M3, M6, M12; Figure 2B; Supplementary Table S2; Supplementary Table S3). By contrast, the modules showing increasing expression across embryogenesis and later stages tended to have significant enrichment of developmental GO and behavioral PEA terms, such as regulation of cell shape, neural activity, linker cell migration, and animal motility (Figure 2B, purple and green shaded modules; Supplementary Table S2; Supplementary Table S3). The most overrepresented terms across all co-expression modules were found in early-embryogenesis module M4, involving 21-fold enrichment of genes associated with protein heterodimerization activity (GO) and 19-fold enrichment of early embryonic chromatid segregation (PEA) (Figure 2B; Supplementary Table S2; Supplementary Table S3). Among the 105 genes in the *C. elegans* genome annotated with the protein heterodimerization activity GO term (GO:0046982), 69% correspond to histones, with most of the others comprised of TATA-box binding proteins, transcription factors, and CENP centromere-related proteins; M4 alone has 31 histones.

### Rapid molecular evolution of genes with peak expression in early embryogenesis and adulthood

The co-expression modules differ significantly in the rate at which their gene members evolve (n=12,628 genes with both expression and divergence information; Figure 3). Surprisingly, we found that it is those genes in M4 with peak expression in early embryogenesis that comprise the most rapidly-evolving set of genes (median *K*_A_ = 0.43; Figure 3). As another sign of rapid evolution of genes in M4, this module contained the lowest percentage of genes with identifiable orthologs between *C. elegans* and *C. briggsae* (28% vs. 64% genome-wide and 92% ortholog pairs identified for M6; Figure 3). The saturated synonymous-site divergence for *C. elegans* orthologs precludes robust tests of adaptive evolution (median *K*_S_ = 2.33), though a large fraction (83%) of non-synonymous substitutions are estimated to have been driven by positive selection in other *Caenorhabditis* (Galtier 2016).

Curiously, however, module M4 has the highest fraction of genes with near-zero values of *K*_A_ (9.3% vs. 0.5% of genes overall; Supplementary Figure S3). This observation indicates exceptionally strong selective constraint on this subset of genes within M4: this subset is comprised entirely of histones which are well-known to evolve slowly, and yet are still overrepresented in M4. These 14 histone genes, plus another subgroup of 15 genes with *K*_A_ < 0.02 (14 of which also are histones), imply that about 20% of M4’s “early embryogenesis” genes encode histones, genes that evolve extraordinarily slowly. Nevertheless, the remaining 80% evolve so remarkably fast that they confer on M4 the highest average *K*_A_ of any module (Figure 3; Supplementary Figure S3). The only other module with substantial abundance of a group of exceptionally conserved coding sequences is ‘constitutive’ module M6 (4.9% of genes with near-zero *K*_A_), which also shows the strongest sequence conservation on average irrespective of this exceptional subset of genes. Module M6 has a median *K*_A_ = 0.033, implying that only about 3% of non-synonymous sites in codons have changed between *C. elegans* and *C. briggsae* since their common ancestor, estimated at 113 million generations ago (Cutter 2008).

The four modules with peak post-embryonic expression and enrichment with spermatogenic function also evolve up to twice as rapidly as the genome-wide median *K*_A_ = 0.121 (median *K*_A_ for “post-embryonic” modules M9, M10, M11, M14 from 0.185 to 0.261; Figure 3). Overall, co-expression modules with lower incidence of sex-neutral genes exhibit more rapid sequence divergence (Figure 4). As expected from previous analyses of *C. elegans* molecular evolution (Cutter et al. 2009), genes in those modules with higher average expression tend to evolve more slowly and show more sequence conservation (Figure 4); this manifests as unusually low divergence at synonymous sites only for M6 (median *K*_S_ = 1.1 vs. genome-wide median *K*_S_ = 2.33). An outlier to the *K*_A_-expression relationship, however, is module M4: these early-embryogenesis genes show fast molecular evolution despite relatively high transcript levels (Figure 4). Our gene-wise analysis of coarse-grained cubic function parameters corroborate these findings (Supplementary Figure S4), with the four α and β parameters being capable of explaining 11.5% of the variability in *K*_A_ across genes (ANOVA F_4,12623_ = 408.5, P<0.0001; log-transformed *K*_A_).

**Figure 4.**
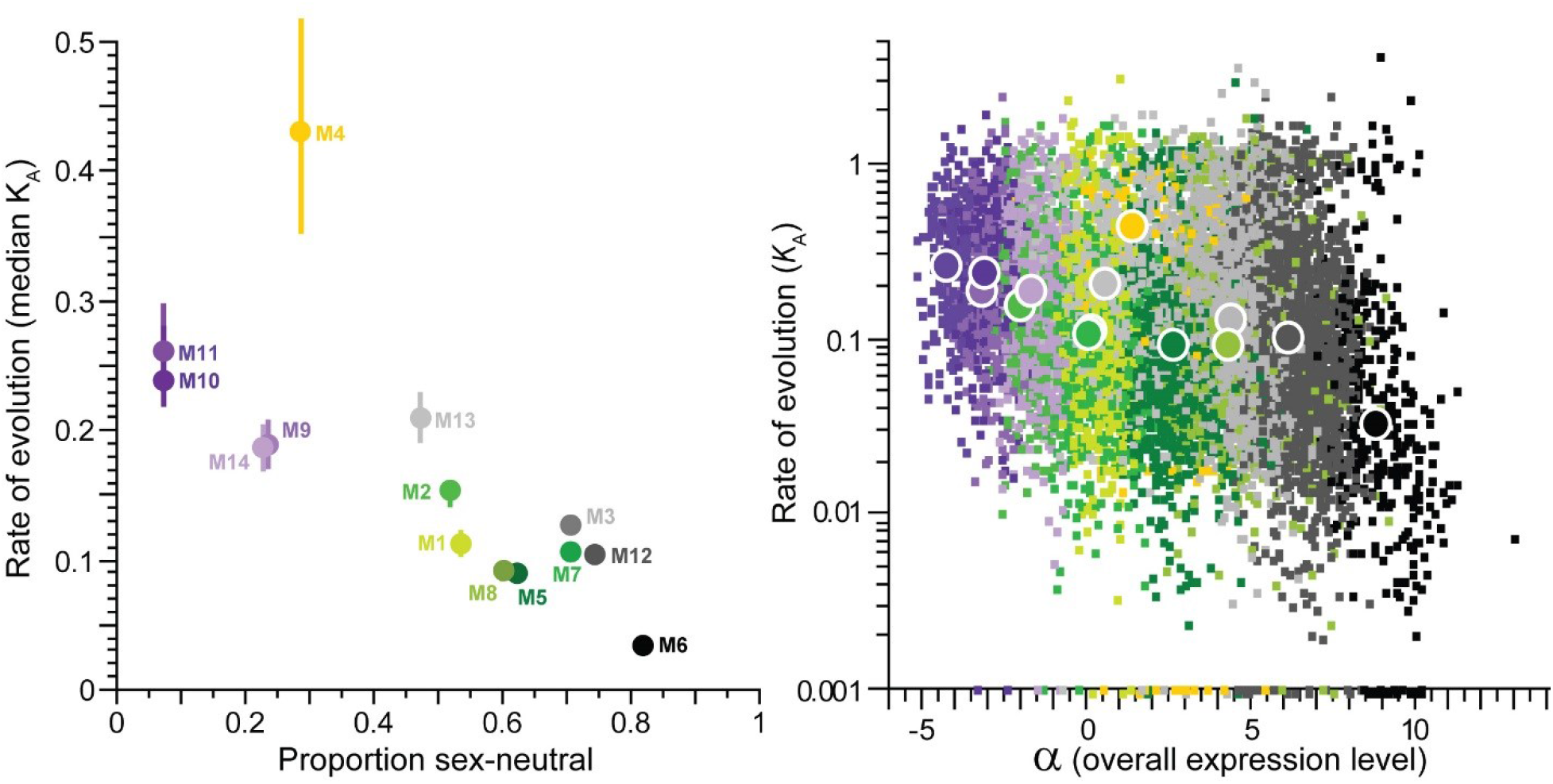
(A) Median rate of protein evolution (non-synonymous site substitution, *K*_A_ ± interquartile range for orthologs of *C. elegans* and *C. briggsae*) for genes within each co-expression module as a function of the proportion of module genes with the sex-neutral expression category, as defined by Ortiz et al. (2014). (B) Rates of protein evolution (*K*_A_, log-scale; zero values plotted at *K*_A_ = 0.001) plotted as a function of the α parameter (overall expression level) from the polynomial fit to the expression time series. Per-gene values shown as small squares, module median values shown as large circles. Module membership color is the same in A and B.

As a complement to the ontogenetic expression module analysis, we quantified rates of molecular evolution for a simpler partitioning of genes, by grouping genes according to the timepoint with highest observed expression. Average rates of protein sequence evolution were fastest for those genes with peak expression in the final L4 larval stage, young adults and in early embryos (Figure 5), corroborating the findings from the ontogenetic co-expression modules. Among those genes with peak expression in embryogenesis, genes with later peak expression tended to evolve more slowly (Figure 5), recapitulating the contrast of *K*_A_ for “early embryogenesis” module M4 versus “late embryogenesis” co-expression modules (M1, M2, M5, M7, M8).

**Figure 5.**
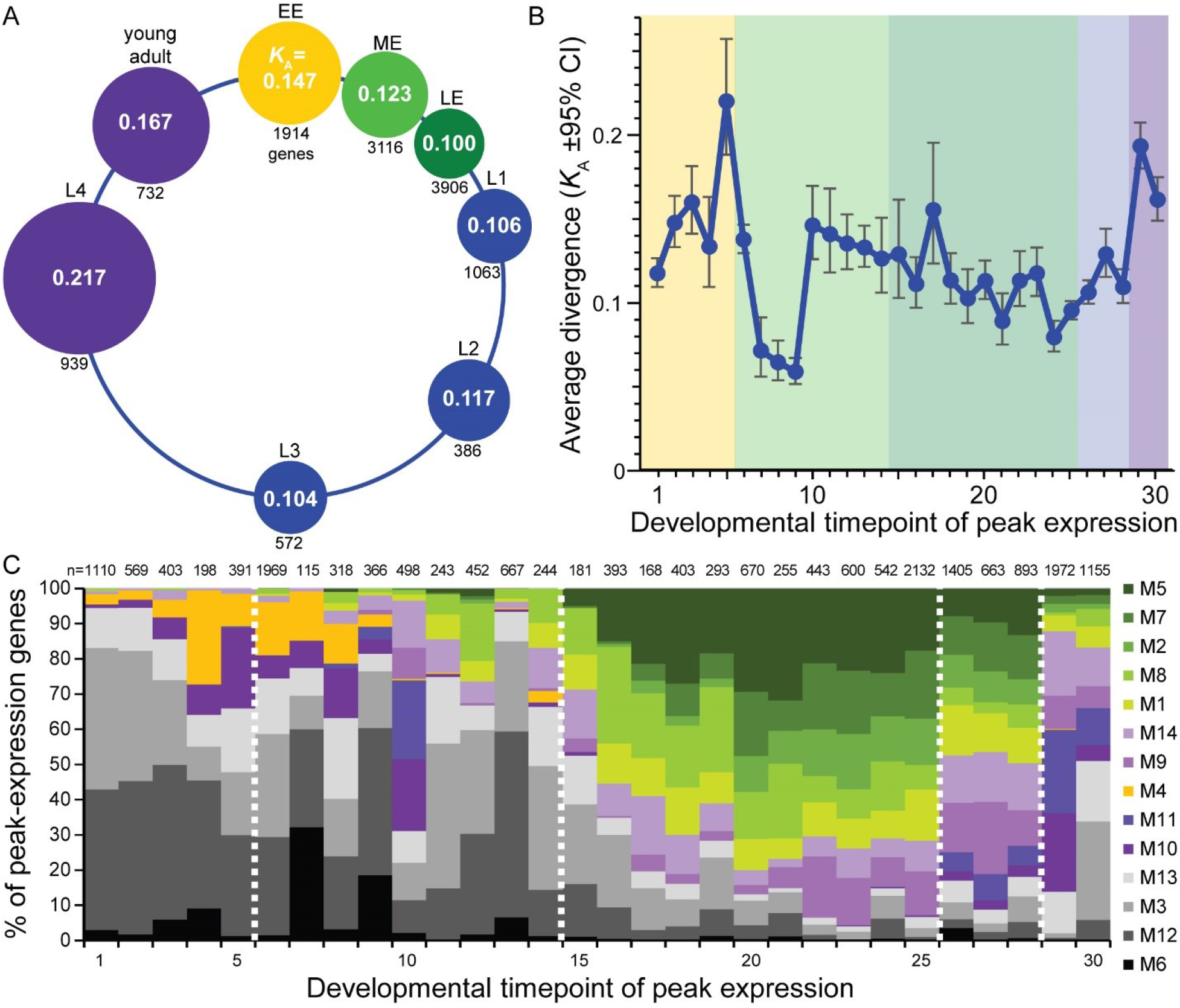
(A) Median rates of protein evolution (*K*_A_) for genes with peak expression in different stages of the *C. elegans* life cycle (EE = early embryo to 120 min, ME = mid-embryo to 390 min, LE = late embryo to 720 min, larval stages L1-L4, young adult); circle diameters proportional to value of *K*_A_. (B) Average *K*_A_ for genes with peak expression at each timepoint in the ontogenetic time series (from back-transform of mean log values; shading: yellow = EE, light green = ME, dark green = LE, blue = larval L1-L3, purple = larval L4 to young adult). Timepoints in embryogenesis are spaced at 30 minute intervals (Gerstein et al. 2010; Gerstein et al. 2014). (C) Cumulative fraction of genes having peak expression at each timepoint that are members of the 14 co-expression modules (number of genes with peak expression at a given timepoint indicated above each bar). Module identities sorted by expression profile similarity as in Figure 1 and colored as in Figure 4; dashed vertical white lines demarcate the boundaries between EE, ME, LE, larval L1-L3, larval L4 to adult as in B.

Interestingly, however, genes with peak expression at timepoints 7-9 (180-240 minutes) exhibit a dip in sequence divergence (Figure 5), suggesting a trend of greater sequence conservation near ventral enclosure in embryogenesis reminiscent of “hourglass” patterns of expression divergence between species (Levin et al. 2012). Caveats to concluding that this observation strongly supports an “hourglass” model of sequence evolution include the facts that timepoints 7-9 exhibit among the fewest genes with peak expression (from 115 genes in timepoint 7 to 366 in timepoint 9) and the clustering analysis revealed no distinct co-expression module exhibiting maximal expression in this developmental interval. Moreover, genes in the highly conserved and highly expressed “constitutive” modules M6 and M12 predominate among the genes with nominally peak expression between timepoints 7-9 (Figure 5), with histone genes especially enriched in timepoint 8. To test the sensitivity of these results to gene sample size and composition, we calculated the “transcriptome divergence index” (TDI; (Quint et al. 2012)), a metric of average sequence evolution for all 12,628 genes with *K*_A_ values weighted by their expression level at a given timepoint (Supplemental Figure S5). Inspection of TDI over the time series shows TDI minimized at timepoint 7 (Supplemental Figure S5), suggesting that ventral enclosure may indeed represent a crucial developmental stage in terms of both conservation of the expression and sequence of genes (Levin et al. 2012). The TDI metric also is especially low at timepoint 1, perhaps consistent with the “early conservation” model, though these earliest transcripts likely are primarily maternal in origin. TDI has a maximal value in adulthood (Supplemental Figure S5), also showing high values in early embryonic developmental timepoints 2-6, consistent with our observations for ontogenetic co-expression modules and the peak expression analysis.

## Discussion

Understanding the interplay between genes and phenotypes in the evolution of development must accommodate how molecular evolution can associate with both phenotypic divergence and phenotypic conservation. The conservation of phenotype, including developmentally static phenotypes like *Caenorhabditis* embryogenesis (Zhao et al. 2008; Memar et al. 2018), need not imply conservation of the genetic pathways that produce them (Kalinka and Tomancak 2012). This idea is the essence of developmental system drift (DSD) (True and Haag 2001), and a key question is to what extent are different stages of development more or less susceptible to molecular divergence and DSD in a predictable way. Temporal trajectories of gene co-expression provide a means of interrogating this question to determine what are the rules in the molecular evolution of development.

### Timing and breadth of expression in the molecular evolution of development

We observe the fastest coding sequence evolution for genes with peak expression early in embryogenesis (co-expression module M4), suggesting that this developmental stage in *C. elegans* near gastrulation may be especially prone to DSD. Why do genes with peak expression in early embryogenesis evolve so fast? This rapid evolution occurs despite an over-representation of histone proteins within this co-expression module that have exceptionally slow sequence evolution. Among the genes with rapid evolution, weaker purifying selection on maternally-provisioned transcripts provides one plausible basis for faster evolution of early-embryogenesis genes (Cruickshank and Wade 2008). A greater incidence of positive selection also could contribute to the rapid evolution of genes in M4, perhaps resulting from parent-offspring conflict or protein-protein co-evolution yielding DSD between gene partners (True and Haag 2001; Clark et al. 2009; de Juan et al. 2013). Moreover, genes in M4 are over-represented on autosomal arms (64% of M4 genes on arms vs. 37% genome average), genomic regions known to harbor genes with greater divergence (Cutter et al. 2009). Despite the extreme consistency of cell lineage in early embryos of different *Caenorhabditis* species (Zhao et al. 2008; Memar et al. 2018), the underlying molecular controls of early embryogenesis have diverged radically so that embryonic arrest near this stage represents the usual fate of inter-species hybrids (Baird et al. 1992; Baird and Seibert 2013; Dey et al. 2014; Bundus et al. 2015), consistent with divergence of genetic interactions with important biological consequences. Thus, the molecular evolutionary consequences of the biased composition of genes with peak expression early in embryogenesis might be predisposed to DSD and to contribute to hybrid inviability in the speciation process.

Our observation of more rapid coding sequence evolution for genes with peak expression early in embryogenesis clearly conflicts with the “early conservation” model for the evolution of development (Kalinka and Tomancak 2012). Moreover, it has been argued that “conservation at the end of embryogenesis is not endorsed by any model” (Kalinka and Tomancak 2012), and yet the trend we observe shows just that, based on analyses of both co-expression modules and peak gene expression patterns. Our analysis of peak expression timing and an expression-weighted divergence index (Figure 5B, Supplementary Figure S5), however, hint at a phase of mid-embryonic development with strongest constraint (timepoint 7, at 180 min), suggestive of the “hourglass model” that has been endorsed in *Caenorhabditis* from analysis of expression divergence (Levin et al. 2012; Levin et al. 2016). The prevalence of genes from highly expressed “constitutive” co-expression modules during the “waist” of the hourglass, however, makes it challenging to understand what is distinctive about the genes with expression at this point midway through embryogenesis. Possible factors could involve the abundance of histone genes to define it as a key developmental phase for chromatin remodeling; alternately, this timepoint might simply represent a lull in stage-specific expression with the “constitutive” genes inevitably dominating the expression composition and, consequently, the signal of high sequence conservation. Regardless, the unusually rapid evolution of early embryogenesis represents, in our view, the pattern of molecular evolution requiring special explanation and attention. Developmental stages associated with genes having faster rates of molecular evolution ought to be predisposed to more extensive developmental system drift, and impose detectable and predictable phenotypic rules. Specifically, stages prone to DSD may reveal themselves by manifesting as being most sensitive to hybrid dysfunction in crosses between diverged species (Bundus et al. 2015).

To date, analyses of molecular evolution have primarily revealed gametic and post-embryonic stages to have fastest rates of evolution in animals and plants (Cutter and Ward 2005; Ellegren and Parsch 2007; Arunkumar et al. 2013; Piasecka et al. 2013; Liu and Robinson-Rechavi 2018a). Our findings corroborate this result, showing that co-expression modules with peaks in adulthood that are enriched for sperm-related gene function evolve especially rapidly. In the context of *C. elegans* biology, where self-fertilizing hermaphrodites evolved from an outbreeding male-female species, both sexual selection pressures in the ancestral species and relaxed sexual selection in the modern day likely contribute to the rapid evolution of sperm genes (Cutter et al. 2019).

Tissue-specific genes have faster coding sequence evolution in mammals (Liao and Zhang 2006b), and temporal specificity might lead to similar consequences. In our analysis, we can think of genes with extreme values of β1, β2, and β3 as having greater temporal specificity of expression and therefore mutations to them having lower potential for pleiotropic effects; however, we observe relatively weak individual associations of these metrics with *K*_A_ (Supplementary Figure S2). Alternately, we can think of mutations to genes with lower α (i.e. a profile of lower overall expression across ontogeny) as having lower potential for pleiotropic effects due to the rarity of gene products, and indeed genes with lower α evolve faster. Genes in module M4, with peak expression during early embryogenesis, represent important outliers to this trend, as they tend to have both fast sequence evolution and moderately high values of α (Supplementary Figure S2). In yeast, however, factors like translational robustness appear to be especially important in mediating the correspondence between expression level and rate of coding sequence evolution (Drummond et al. 2005), though it remains unclear how general this explanation holds across eukaryotes.

### Linking divergence in expression with divergence in sequence

Our analysis puts to the side the question of the relative importance of regulatory versus coding changes in adaptation and morphological divergence (Wray 2007; Carroll 2008; Stern and Orgogozo 2008). Instead, we focus on coding sequence evolution to ask what features of ontogeny predict differences in the rates of evolution across genes. However, observing differences in rates of coding sequence evolution among distinct co-expression modules implies a mapping between the nature of regulatory control and protein evolution. Previous studies of diverse animals show a weakly positive correlation between molecular evolutionary rates of coding sequences and regulatory regions (Jordan et al. 2005; Lemos et al. 2005; Liao and Zhang 2006a), including for *Caenorhabditis* (Castillo-Davis et al. 2004; Mark et al. 2019). Both coding sequences and gene expression are subject to purifying selection in *C. elegans* (Denver et al. 2005; Cutter et al. 2009), but future genome-scale analyses that couple ontogenetic transcriptome profiles with coding and regulatory sequence evolution are required to more fully determine the magnitude of inter-dependence of these modes of molecular evolution across development. Establishing such links would be valuable in integrating “hourglass” patterns of expression divergence and sequence evolution.

Evo-devo generally focuses on how the relative strength of constraint, which manifests as purifying selection and sequence conservation, could shape temporal ontogenetic patterns of evolution (Kalinka and Tomancak 2012). And yet, micro-evolutionary studies demonstrate that a majority of amino acid substitutions in protein coding sequence evolution often accumulate as a result of adaptive evolution in many animals, especially those with large effective population sizes like *C. elegans’* congeners (Galtier 2016). Genes biased toward expression in adults and gametes are known to show elevated rates of adaptive evolution (Swanson and Vacquier 2002; Arunkumar et al. 2013; Liu and Robinson-Rechavi 2018a), but the extent of embryonic adaptive evolution and its implications are less well established. In *Drosophila,* rapidly-evolving proteins involved in chromatin regulation and genomic conflict are known to play important roles in creating post-zygotic reproductive barriers between species during early development (Presgraves 2010; Maheshwari and Barbash 2011; Cooper et al. 2018). Evolutionary conflict over allelic expression in early embryos also can drive rapid sequence evolution (Haig 1997). Presuming a substantial contribution of adaptive divergence to coding sequence evolution in *Caenorhabditis* (Galtier 2016), our findings support the possibility that adaptive evolution, rather than weaker constraint, contributes importantly to ontogenetic patterns in the molecular evolution of development (Kalinka and Tomancak 2012). Rapid evolution of genes expressed at distinct times in embryogenesis, whether due to adaptation or weaker constraint, should lead to predictable developmental manifestations in the form of hybrid dysfunction in the speciation process.

## Acknowledgements

We are grateful to the ModENCODE consortium for providing publicly available expression data for *C. elegans.* ADC is supported with funds from the Natural Sciences and Engineering Research Council (NSERC) of Canada; LS is supported with funds from NSERC and the Canadian Institutes of Health Research (CIHR). RG was supported by an NSERC Undergraduate Summer Research Award.

## Author Contributions

ADC, RG and LS conceived and designed the study; ADC, RG, SM, and WW analyzed data; ADC drafted the initial version of the manuscript and all authors contributed to the final manuscript.

## Data Accessibility

Expression data collected by ModENCODE are publicly available at (http://data.modencode.org).

