## Supplementary Tables and Figures for "Molecular evolution across developmental time reveals rapid divergence in early embryogenesis" (from another version of this preprint: version 1, 2019-01-11)

### Supplementary Information

**Supplementary Table S1**. List of ModENCODE *C. elegans* transcriptome datasets used in analysis of differential expression across development.

| ModENCODE ID | Dataset | Stage or embryo age treatment (h) |
|---|---|---|
| 4527 | L1 (4h post-L1) | L1 |
| 4574 | L1 (4h postL1) | L1 |
| 4530 | L2 (17.75h post-L1) | L2 |
| 4575 | L2 (17.75h post-L1) | L2 |
| 4532 | L3 (26.75h post-L1) | L3 |
| 4576 | L3 (26.75h post-L1) | L3 |
| 4534 | L4 (34.25h post-L1) | L4 |
| 4535 | L4 (34.25h post-L1) | L4 |
| 4581 | Young adult (pre-gravid, 46h post-L1) | adult |
| 4593 | Young adult (pre-gravid, 46h post-L1) | adult |
| 6561 | N2_EE_50-0 | 0 |
| 6568 | N2_EE_50-30 | 30 |
| 6579 | N2_EE_50-60 | 60 |
| 6585 | N2_EE_50-90 | 90 |
| 6562 | N2_EE_50-120 | 120 |
| 6563 | N2_EE_50-150 | 150 |
| 6564 | N2_EE_50-180 | 180 |
| 6565 | N2_EE_50-210 | 210 |
| 6566 | N2_EE_50-240 | 240 |
| 6567 | N2_EE_50-270 | 270 |
| 6569 | N2_EE_50-300 | 300 |
| 6570 | N2_EE_50-330 | 330 |
| 6571 | N2_EE_50-360 | 360 |
| 6572 | N2_EE_50-390 | 390 |
| 6573 | N2_EE_50-420 | 420 |
| 6574 | N2_EE_50-450 | 450 |
| 6575 | N2_EE_50-480 | 480 |
| 6576 | N2_EE_50-510 | 510 |
| 6577 | N2_EE_50-540 | 540 |
| 6578 | N2_EE_50-570 | 570 |
| 6580 | N2_EE_50-600 | 600 |
| 6581 | N2_EE_50-630 | 630 |
| 6582 | N2_EE_50-660 | 660 |
| 6583 | N2_EE_50-690 | 690 |
| 6584 | N2_EE_50-720 | 720 |

**Supplementary Table S2**. Summary of gene ontology (GO) term enrichment for each co-expression module.

**Supplementary Table S3**. Summary of phenotype enrichment analysis (PEA) terms for each co-expression module.

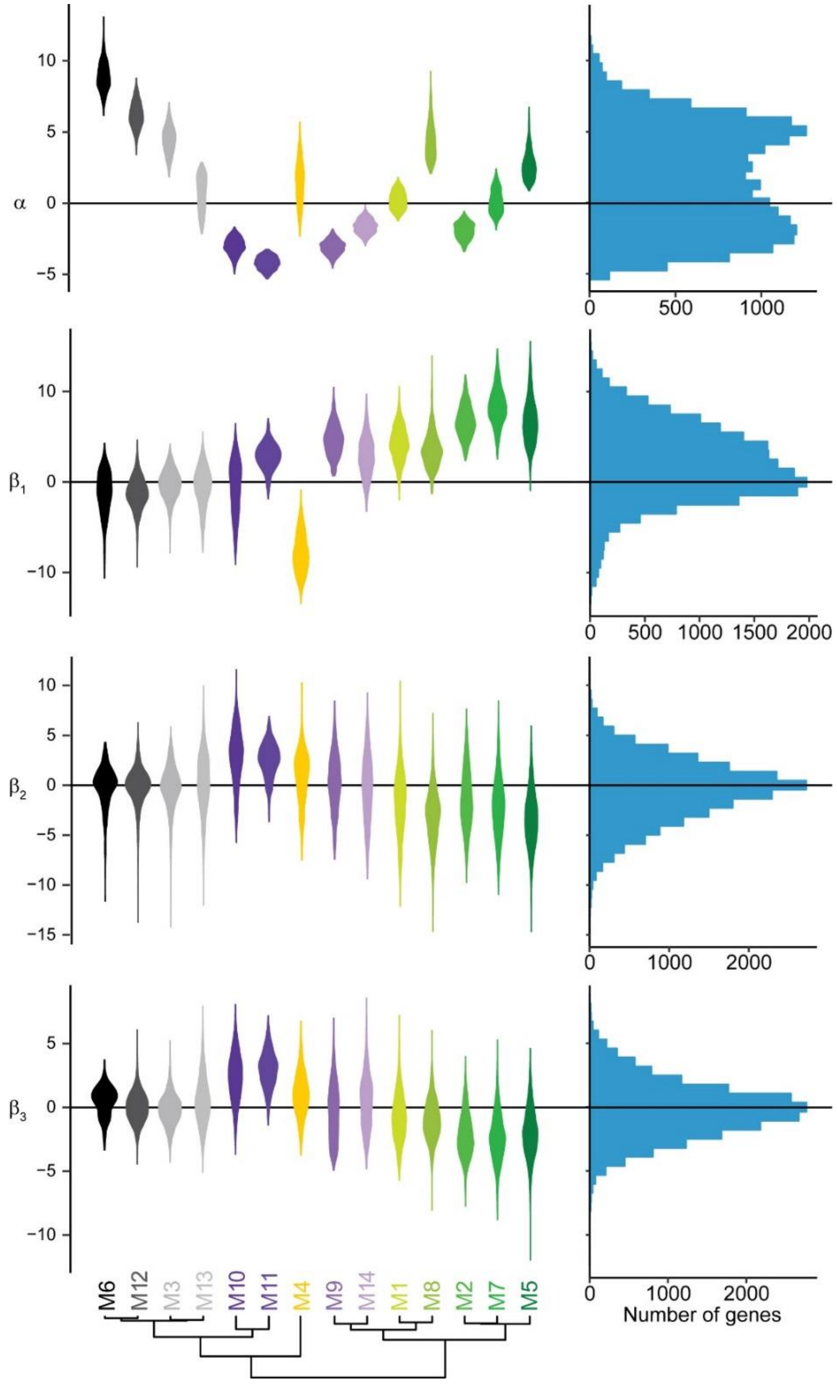


**Figure S1**. Distributions of polynomial parameter values from expression profile function fits for each gene ($\alpha$ = overall expression level, $\beta_1$ = linear change over time, $\beta_2$ = quadratic curvature, $\beta_3$ = cubic S-shape to expression profile over development). Violin plots show the parameter value densities separately for each co-expression module (see Figure 1 in main text).

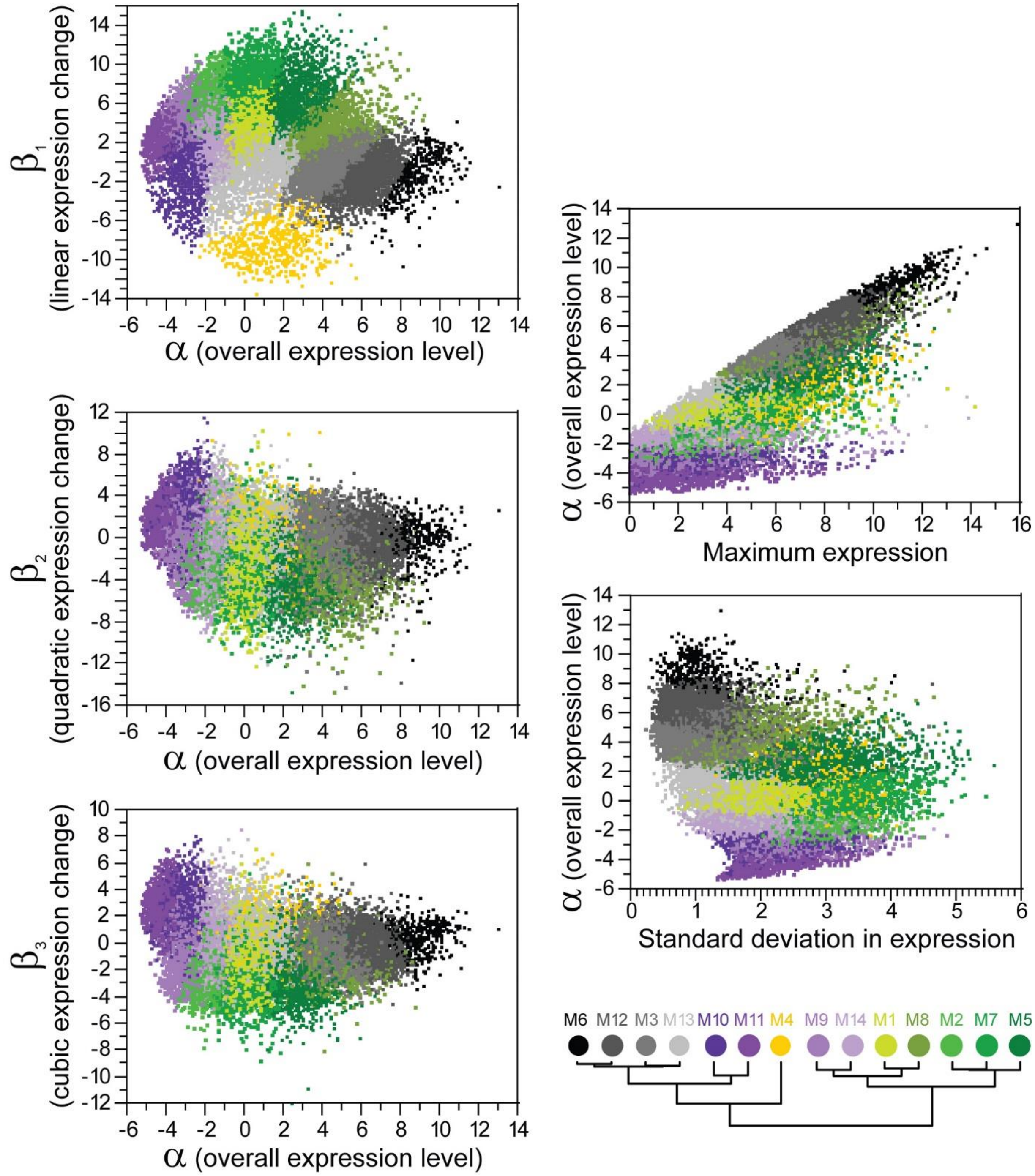


**Supplementary Figure S2**. Biplots of per-gene polynomial fit parameter values, colored by co-expression module (as in Figure 4 main text), show clustering of genes with similar parameter values ($\alpha$ = overall expression level, $\beta_1$ = linear change over time, $\beta_2$ = quadratic curvature, $\beta_3$ = cubic S-shape to expression profile over development). The $\alpha$ parameter correlates positively with a measure of maximum expression level across timepoints and has a complex relationship with variation among timepoints in expression level.

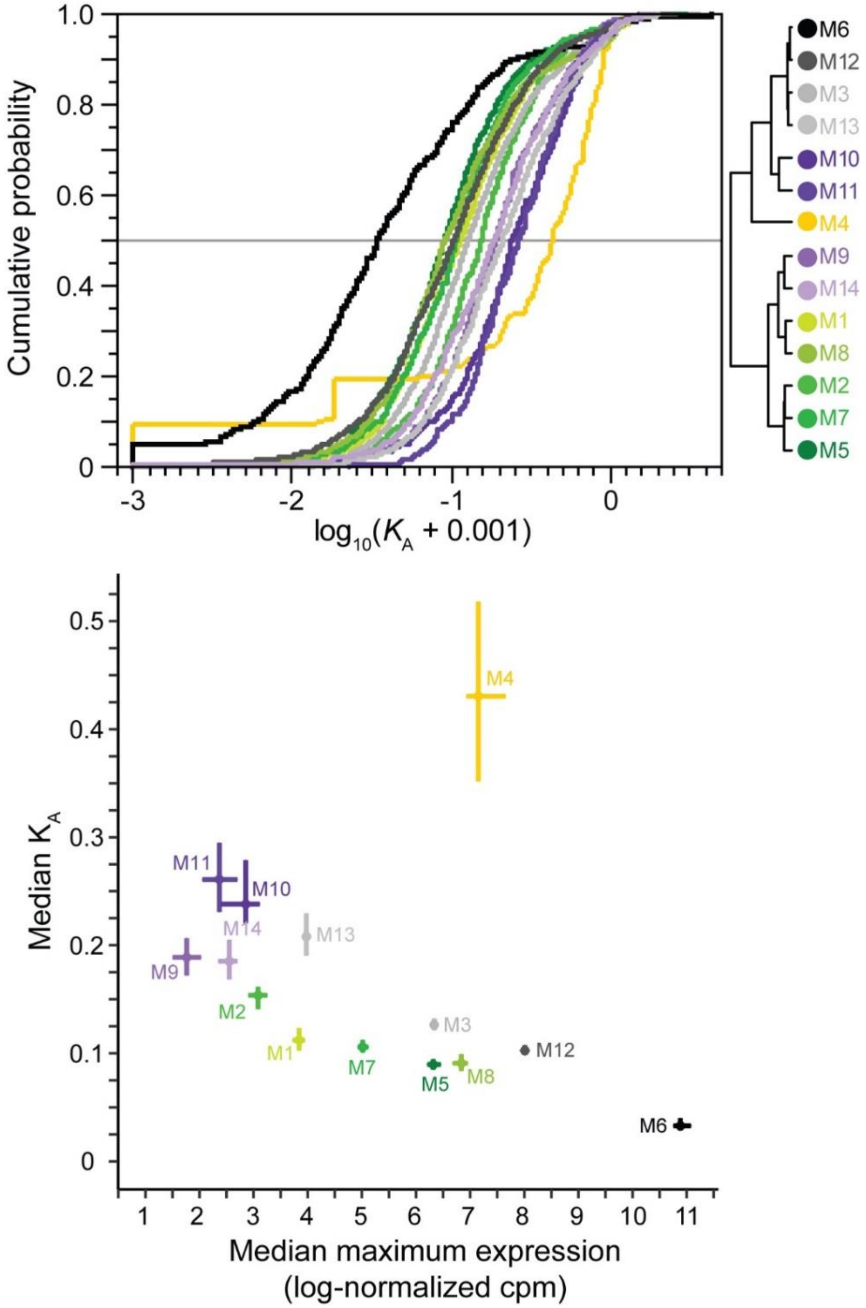


**Supplementary Figure S3**. Cumulative distribution of non-synonymous site substitutions (log-transformed $K_{\mathrm{A}}$) for each co-expression module illustrates the distinct incidence of extremely low $K_{\mathrm{A}}$ values for M4 and M6 (top panel), indicating the subset of genes with little protein sequence divergence between *C. elegans* and *C. briggsae*. Otherwise, genes in M6 have a low median rate of evolution whereas genes in M4 have a high median rate of evolution (horizontal line indicates 50$^{\mathrm{th}}$ percentile). Modules containing genes with higher average expression tend to have slower rates of evolution (bottom panel); note M4 as outlier to this trend.

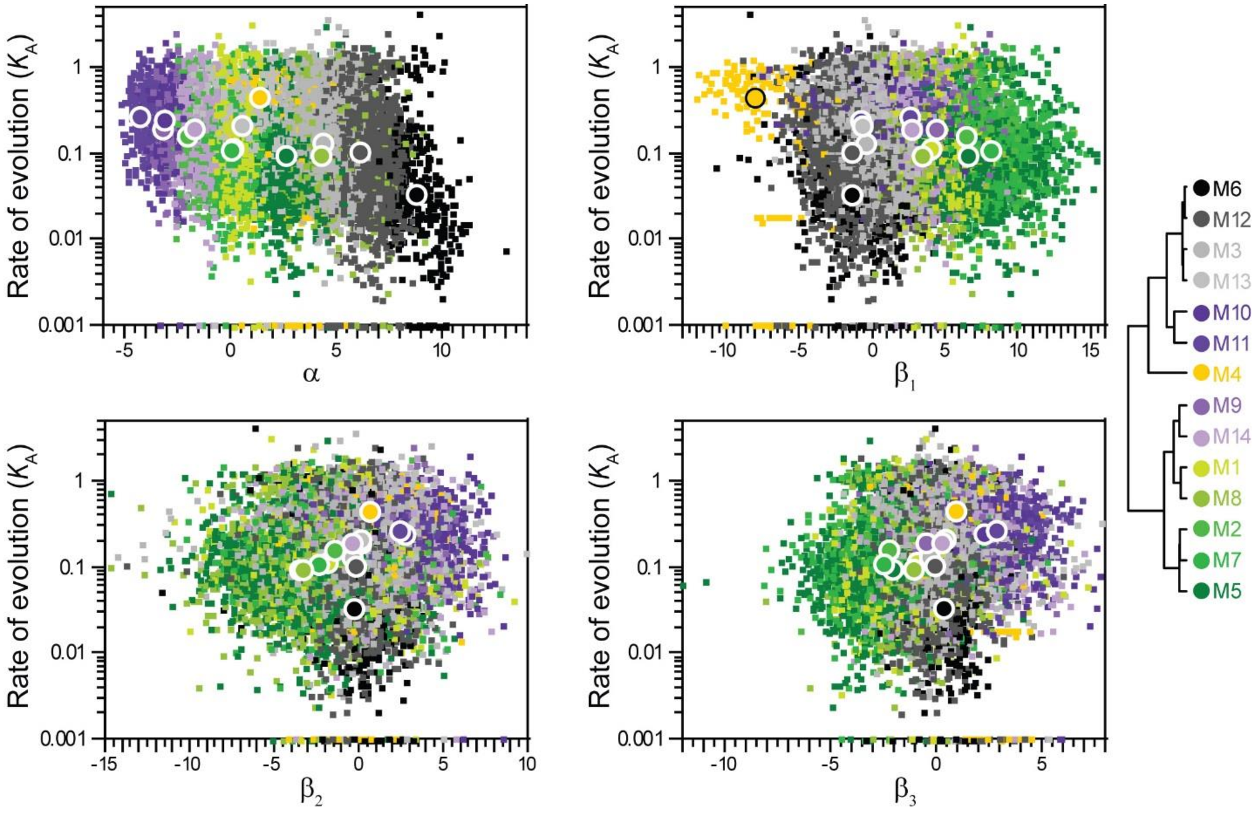


**Supplementary Figure S4**. Rates of protein evolution ($K_A$, log-scale) plotted as a function of the polynomial fit parameter values to the expression time series ($\alpha$ = overall expression level, $\beta_1$ = linear change over time, $\beta_2$ = quadratic curvature, $\beta_3$ = cubic S-shape to expression profile over development). Per-gene values shown as small squares, module median values shown as large circles. Dendrogram indicates expression similarity among co-expression modules.
